# Determining the Hierarchical Architecture of the Human Brain Using Subject-Level Clustering of Functional Networks

**DOI:** 10.1101/350462

**Authors:** Teddy J. Akiki, Chadi G. Abdallah

## Abstract

Optimal integration and segregation of neuronal connections are necessary for efficient large-scale network communication between distributed cortical regions while allowing for modular specialization. This dynamic in the cortex is enabled at the network mesoscale—the organization of nodes into communities. Previous *in vivo* efforts to map the mesoscale architecture in humans had several limitations. Here we characterize a consensus multiscale community organization of the functional cortical network. We derive this consensus from the clustering of subject-level networks. We show that this subject-derived consensus framework yields clusters that better map to the individual, compared to the widely-used group-derived consensus approach. We applied this analysis to magnetic resonance imaging data from 1003 healthy individuals part of the Human Connectome Project. The hierarchical atlas and code will be made publicly available for future investigators.

## 1. Introduction

Using surrogates of brain activity such as the blood-oxygen-level dependent (BOLD) signal obtained using functional magnetic resonance imaging (fMRI), whole-brain functional networks (i.e., *connectomes*) can be estimated *in vivo*. The brain functional connectome is organized at multiple spatial scales, one of which is the *mesoscale*. It has been recently shown that a full repertoire of functional communities—groups of nodes that are densely connected internally—can be consistently decoded, even at rest. In the brain, these communities are known to represent subsystems and mediate distinct neurophysiological functions (e.g., the brain’s visual subnetwork) (Yeo et al., 2011; Power et al., 2011; Cole et al., 2014). Moreover, this scale is highly sensitive to disease, where several psychiatric disorders have shown selective disruption in particular brain communities (Alexander-Bloch et al., 2012; Akiki et al., 2017, 2018; Menon, 2011; Cisler et al., 2016). This apparent importance has prompted interest in this line of investigation; while a vast wealth of knowledge has been gained from these efforts, several methodological pitfalls remain.

To map the mesoscale architecture of the normal brain, previous studies have generally applied the community detection algorithm on a group-representative network obtained by averaging networks from a group of individuals. However, it is becoming increasingly clear that important features may be lost by such averaging (including some that are present across individuals (Gordon et al., 2017a)), leading to a representation that may not resemble a true central tendency in the group (Dubois and Adolphs, 2016; Hacker et al., 2013; Gordon et al., 2017b; Braga and Buckner, 2017; Finn et al., 2015; Laumann et al., 2015).

Further, the analysis of networks derived from time series (i.e., correlation networks) is challenging. First, unlike prototypical networks where edges are either present or absent (Euler, 1736), edges in correlation networks represent the magnitude of statistical association, and so are on a continuum. Second, methods that are commonly used to index association— most commonly Pearson correlation—also produces negative values. Third, these networks contain numerous indirect dependencies by virtue of the transitivity inherent in correlations (Zalesky et al., 2012; Barzel and Barabási, 2013; Giraud et al., 1999). This is particularly salient in large networks, as these indirect effects may be compounded with higher-order interactions.

Previous investigations have generally made use of thresholds prior to analysis in order to treat functional networks like usual sparse graphs. The rationale is that the strong connections that would be retained would likely be the most relevant and least likely to be artifactual, also leading to the elimination of negative edges. Recently, the relevance of weak network edges has become increasingly recognized, as they appear to convey unique information not encoded in strong edges (Santarnecchi et al., 2014; Lohse et al., 2014; Bassett et al., 2012; Gallos et al., 2012; Goulas et al., 2014). Further, the choice of the threshold is also challenging and can give rise to heterogeneity in the findings (Jalili, 2016; Garrison et al., 2015). Like weak edges, negative edges (i.e., anticorrelations) have been also shown to have a substantial physiological basis (Nielsen et al., 2018; Fox et al., 2005; Zhan et al., 2017; Parente et al., 2017; Kelly et al., 2008).

While several methods exist for community detection in networks, most are not compatible with weighted networks containing both positive and negative links of the type observed in correlation networks (Fortunato, 2010; Sporns and Betzel, 2016). One of the most commonly used and best-performing methods in community detection is the Louvain algorithm (Blondel et al., 2008; Lancichinetti and Fortunato, 2009; Sporns and Betzel, 2016). The Louvain algorithm is versatile and can be used with different null models, some of which have been extended to allow for positive and negative interactions (Sporns and Betzel, 2016). Importantly, null models that are typically used with the Louvain algorithm are based on permutations (rewiring) of the original networks, preserving the total weight and degree while randomizing connections (Girvan and Newman, 2002). This is known as the Newman-Girvan (NG) null model, which has also been extended to signed networks (Gómez et al., 2009; Rubinov and Sporns, 2011; Traag and Bruggeman, 2009). However, such an approach is problematic in the case of correlation matrices, as it assumes that the entries are independent, thereby violating the correlation transitivity (Bazzi et al., 2016; MacMahon and Garlaschelli, 2015). Therefore, the community detection may not be accurate, particularly if there is heterogeneity in the size of the communities (Bazzi et al., 2016)—which is known to be biologically plausible in the case of the brain.

A number of recent developments in network science and neuroimaging have paved the way for frameworks that can be used to address these challenges. Here we used the Louvain algorithm, with a null model based on random matrix theory designed explicitly for correlation networks (we refer to it throughout as the RMT null model) (MacMahon and Garlaschelli, 2015). While it was initially introduced in finance (MacMahon and Garlaschelli, 2015), it has been recently successfully applied to neurophysiological recordings (Almog et al., 2017). This method, which does not violate transitivity, is compatible with weighted and signed networks, forgoing the need to perform any thresholding. The introduction of more efficient algorithms, of reclustering frameworks, and the increased availability of high-performance computing, have made it practical to perform the clustering at the level of the individual network. As a neuroimaging dataset, we used a recent release from the Human Connectome Project (S1200 release; March 2017) (Essen et al., 2013). This relatively-large sample of healthy individuals consists of state-of-the-art MRI data and addresses several of the limitations present in older datasets (Uğurbil et al., 2013; Glasser et al., 2016b).

We hypothesized that determining the mesoscale architecture of the average human cortex can be better achieved by first mapping the mesoscale in each subject and then arriving at a meaningful central tendency, rather than by mapping the mesoscale of an averaged brain network. To this aim, we perform the clustering at the subject level followed by a reclustering procedure to reach a population consensus mesoscale architecture (we refer to this as *subject-derived consensus*) (Jeub et al., 2018; Lancichinetti and Fortunato, 2012). We compare this subject clustering-reclustering framework to the traditional group-representative method (we refer to it as *group-derived consensus*) to assess which results in solutions that map best to the individuals’ data.

## 2. Materials and Methods

### 2.1. Neuroimaging dataset

The data that we use in this work is from the Washington University-Minnesota Consortium Human Connectome Project (HCP) (Essen et al., 2013). The latest release at the time of writing was adopted (S1200; March 2017). Details regarding this dataset have been previously published (Essen et al., 2013; Glasser et al., 2016b). Briefly, it included state-of-the-art whole-brain MRI acquisition with structural, functional, and diffusion-weighted imaging; the scanner was a customized Siemens Skyra 3T scanner with slice-accelerated sequences for fMRI (Moeller et al., 2010; Feinberg et al., 2010; Setsompop et al., 2011; Smith et al., 2013). Whole-brain functional data were acquired in two sessions. Each session consisted of two phase-encoding directions (left-right and right-left) each a 15 min multiband gradient echo-planar resting-state run (Voxel size = 2.0 × 2.0 × 2.0 mm; TR = 720 ms; TE = 33.1 ms; Flip angle = 52°; 72 slices; Bandwidth = 2,290 Hz/pixel; FOV = 208 × 180 mm). Informed consents were obtained from all subjects. The study procedures were approved by the institutional review boards. This release includes 1097 subjects with resting-state fMRI scans. Our analysis was restricted to subjects who completed all four resting-state scan runs; namely, two sessions, each with two encoding runs. This resulted in n = 1003 individuals; 534 females and 469 males. The age range was 22 to 37, with a mean of 28.7.

### 2.2. First-level processing

The HCP minimal processing pipeline was used (Glasser et al., 2013; Smith et al., 2013). Briefly, this included projection to the surface space, 2 mm FWHM smoothing, ICA+FIX denoising with minimal high-pass filtering, and surface registration using MSMall (Jenkinson et al., 2002, 2012; Fischl, 2012; Robinson et al., 2014, 2018; Griffanti et al., 2014; Salimi-Khorshidi et al., 2014). To define our ROIs, we used a newly-developed multimodal parcellation (MMP) (Glasser et al., 2016a). We chose this parcellation because it is neuroanatomically informed, with data from cortical architecture, connectivity, and topography. This group parcellation consisted of 360 ROIs (180 per hemisphere) that cover the entirety of the cerebral cortex (but does not include the subcortex or cerebellum), which we mapped to the individuals. In the standard surface space, we calculated the mean time series from all voxels in each ROI. Resting-state fMRI time courses from the left-right and right-left phase-encoding runs and the two sessions were concatenated (total duration 60 minutes; 4800 time point). Functional connections between nodes *i* and *j* were defined as the Pearson product-moment correlation of their respective time series.

### 2.3. Community detection

This consisted of using the Louvain community detection algorithm (Blondel et al., 2008); specifically, an iterated version as implemented in the GenLouvain package (Jeub et al., 2017). This versatile method is based on finding communities with a high degree of intra- and low degree of interconnectedness. This is achieved heuristically by optimizing the modularity metric *Q*, which captures the degree of connectedness within communities compared to connectedness expected under a null model (Blondel et al., 2008; Girvan and Newman, 2002):

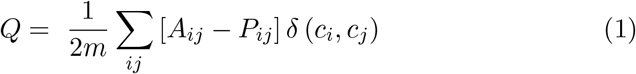

where *A* and *P* are the observed and null adjacency matrices, respectively; and 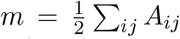 is the total strength of the network. The Kronecker *δ* (*c_i_, C_j_*) function is equal to 1 when *i* and *j* are in the same community and 0 otherwise. As mentioned in the introduction, classical formulation of *P* is that of a permutation null model (Girvan and Newman, 2002): 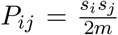 where 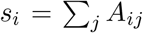 is the strength of node *i*. Although this definition was extended to weighted signed networks (Gómez et al., 2009; Rubinov and Sporns, 2011; Traag and Bruggeman, 2009), it remains inappropriate for use with networks generated from correlation of time series. Namely because entries in the correlation matrices are not independent, which leads to a bias in community detection (Bazzi et al., 2016; MacMahon and Garlaschelli, 2015; Almog et al., 2017; Zalesky et al., 2012; Barzel and Barabási, 2013). We avoided this by adopting a null model that was specifically designed for correlation-based networks (MacMahon and Garlaschelli, 2015) (detailed below). The networks were not thresholded, retaining all weights (both positive and negative).

Methods related to modularity maximization are known to suffer from the resolution limit—the inability to detect communities smaller than a certain size (Fortunato and Barthelemy, 2006). Several methods have been used to address this (Traag et al., 2011; Reichardt and Bornholdt, 2006; Fortunato, 2010; Nicolini et al., 2017). Here we adopt a recursive hierarchical approach to recover the community structure at multiple scales (Jeub et al., 2018; Sales-Pardo et al., 2007). After detecting the first-level (i.e., coarsest) communities in the initial run of the algorithm, we define each detected community as a separate subgraph and run the algorithm recursively on each. This is repeated until no more statistically significant subcommunities can be detected. This iterative procedure allows the resolution of a hierarchically nested structure. Indeed, this is consistent with notions of hierarchical organization in the cortex from microscopic tracer studies in other mammals (Felleman and Essen, 1991; Scannell et al., 1999). We use this hierarchical framework at the subject-level, and a slightly different one at the reclustering stages (see below).

Due to the nearly degenerate outputs of the Louvain algorithm (Good et al., 2010), for each subject-level run, we performed 100 iterations and adopted the partition that is most similar to the ensemble, borrowing the approach from (Traud et al., 2011; Doron et al., 2012; Bassett et al., 2013). The rationale is that similarity to the ensemble can be used as a surrogate for stability, and the most stable partition solution is likely the closest to the ground truth (Sporns and Betzel, 2016; Bassett et al., 2013). To evaluate partition similarity, we used the *z*-score of the Rand coefficient (described below) and calculated the pairwise similarity between all 100 partition solutions, selecting for each subject the partition with the highest mean similarity to the ensemble. Since the community detection procedure was applied recursively in a hierarchical manner, 100 iterations and subsequent selection of the representative partition based on similarity was also applied to each subcommunity. This was done in order to avoid adopting an arbitrary solution from each subject.

### 2.4. A random matrix null model for correlation networks

To generate null models that are appropriate for use with correlation networks, we adopted a method based on random matrix theory (RMT). It is described in full detail in the original publication (MacMahon and Garlaschelli, 2015). Briefly, this method builds null networks using a modified Wishart distribution with the same common trend and noise as the observed network, but without the community structure (sum of the random component and the dominant positive component (MacMahon and Garlaschelli, 2015)). This method does not require thresholding and is compatible with negative values. The end result being that positive interactions are maximally concentrated within modules and negative interactions expelled outside.

For the hierarchical recursive application, after identification of a community c in the initial network, we regenerate a null model from time series of nodes belonging to this community, and the community detection is then applied to this subgraph. One criticism of such recursive procedures relates to the fact that there are no clear stopping rules (Fortunato, 2010). Here, it is important to mention that by virtue of the null model that we employ, only communities that are present in the RMT-filtered networks are detected (MacMahon and Garlaschelli, 2015); therefore, the procedure stops automatically whenever no further “statistically significant” subcommunities can be detected.

### 2.5. Partition similarity

In order to measure the similarity between two partitions, we adopted the *z*-score of the Rand coefficient (Traud et al., 2011; Doron et al., 2012). In the original definition, the Rand coefficient of two partitions is calculated as the ratio of the number of node pairs classified in agreement in both partitions, to the total number of pairs (Rand, 1971). Similar to Doron et al. (2012); Akiki et al. (2018); Betzel et al. (2015); Bassett et al. (2013), here we use the *z*-score variant introduced in Traud et al. (2011), which can be interpreted as a measure of how similar two partitions are, beyond what might arise at random (Traud et al., 2011; Sporns and Betzel, 2016; Fortunato and Hric, 2016). An analytical formula for the z-score of the Rand coefficient between partition *a* and *b* can be written as:

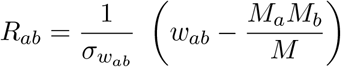

Where *M* is the total number of node pairs, *M_a_* and *M_b_* the numbers of node pairs in the same community in *a* and *b* respectively, *w_ab_* the number of pairs that figure in the same community in both *a* and *b*, and *σ_wab_* the standard deviation of *w_ab_* (as in Traud et al. (2011)). For more information, see SI.

### 2.6. Hierarchical consensus reclustering

Our multiresolution co-classification matrices embed information from the different hierarchical levels of community structures detected from the clustering results at the level of the subjects’ networks. The recently developed method that we adopted extends the classical formulation of consensus reclustering (Lancichinetti and Fortunato, 2012) by allowing a hierarchical multiresolution output and has built-in tests for statistical significance (Jeub et al., 2018). Here, the quality function was modularity-like as in Eq. 1, with *a* “local” variant of the permutation null model. Briefly, in addition to the constraint of the traditional permutation model of a fixed size and number of communities, this local model also assumes that node *i* is fixed and node *j* is random when calculating 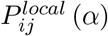. This means that the null network only receives contributions from nodes that are less frequently co-classified together compared to the local permutation model, at a statistical significance level *α*. More detail can be found in Jeub et al. (2018).

The same recursive principle and Louvain implementation described above were used here. To ensure a stable output, 100 iterations are used at each level (cases of output instability are dealt by meta-reclustering as in Lancichinetti and Fortunato (2012); Jeub et al. (2018)); the threshold for statistical significance is set at *α* = 0.05.

### 2.7. Partition task homogeneity

Task fMRI data from the Human Connectome Project span several domains (social cognition, motor, gambling, working memory, language processing, emotional processing, and relational processing) (Barch et al., 2013; Glasser et al., 2016b). Here we used the effect size activation maps over 86 task contrasts (group average over 997 subjects from the S1200 release).

We adopted the framework of module *functional homogeneity* (Schaefer et al., 2017; Kong et al., 2018; Gordon et al., 2017b) to gauge the quality of partition solutions across the hierarchical levels as another metric independent of partition similarity. Nodes within well-defined modules should, in principle, respond in agreement across tasks. That is, modules should show a high degree of homogeneity (e.g., uniformly activated or deactivated when completing a certain task). Similar to Schaefer et al. (2017); Kong et al. (2018), for each task, we quantify homogeneity by calculating the negative of the standard deviation of nodal activation (Cohen’s d) values for each module (the sign was flipped so that *lower*, more negative values indicate lower functional homogeneity, while *greater*, less negative values closer to zero indicate higher homogeneity). Of the 86 task contrasts we excluded redundant ones (e.g., the standard deviation from *Faces – Shapes* contrast is identical to the *Shapes – Faces* contrast), retaining 48 contrasts in total.

However, because we were to compare partitions of different size and numbers, this simple definition would not be sufficient (Gordon et al., 2016; Betzel et al., 2017, 2013). For example, modules with a smaller number of nodes may be more homogeneous. To account for this bias, we standardized it against the null distribution obtained by randomly permuting module assignments 10,000 times (keeping the number and size of the communities constant), and expressed the homogeneity as a *z*-score. For each hierarchy, the *z*-scores were averaged over all modules and then over all task contrasts. This resulted in a summary measure how functionally homogeneous each partition solution is, beyond what is expected by the size and number of modules.

While in certain studies have also used the concept of “connectional homogeneity” to assess partitions (Schaefer et al., 2017; Kong et al., 2018), we believe that not to be adequate for the present study due to the transitivity bias of functional networks that we were set to avoid (Zalesky et al., 2012; Barzel and Barabási, 2013).

### 2.8. Analysis of nodal consistency

To quantify how consistently each nodes is assigned to its community, we calculated the nodal consistency for each partition, here defined as the number of times a node is assigned to the same partition across subjects in a particular hierarchy divided by the total number of times that the node has been classified (i.e., to same or to a different community).

To better understand the variability in consistency across nodes, we calculated the following: 1) the nodal strength as a measure of network *hub-ness* calculated as 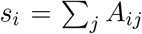 as the strength (hubness) of node *i*, 2) the signal-to-noise ratio defined as 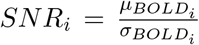 of the nodal BOLD time course, and 3) a measure of nodal activity variation across the task fMRI contrasts, here defined as the coefficient of variation 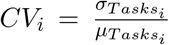. For simplicity, these were calculated from group-averaged data (average connectivity matrix for hubness, group-representative time series for the S1200 sample (Smith et al., 2014) for SNR, and group-average task fMRI contrast maps for the task fMRI coefficient of variation). We then used these terms in a multiple linear regression model as predictors of nodal consistency.

### 2.9. Robustness analyses

In-scanner head motion has been reported to confound the estimation of functional connectivity (Power et al., 2012). For the main analysis, we adopted the strategy proposed by the HCP group (Glasser et al., 2013; Smith et al., 2013) which includes ICA+FIX (Griffanti et al., 2014; Salimi-Khorshidi et al., 2014). We chose not to regress out the “global signal” [mean grey-matter time course regression (MGTR)] as there is concern that the process may distort the correlation structure by shifting the connections to an approximately zero-centered distribution, causing an artifactual increase in computed anticorrelations, and induce a shift in areal boundaries and a distance dependence in functional connections (Murphy et al., 2009; Fox et al., 2009; Glasser et al., 2016a; Power et al., 2014a). The first point is particularly salient in our case as we do not threshold negative connections prior to the community detection analyses. MGTR may therefore spuriously shift (weakly) “positive” corrections into anticorrelations, which would directly impact how they are treated by the RMT-based method, which is based on expelling negative connections outside of modules (MacMahon and Garlaschelli, 2015). However, it has been argued that the standard HCP denoising methods do not fully remove motion artifacts and that regression of the “global signal” remains an effective strategy in reducing the dependence of correlations on motion (Power et al., 2017a, b; Burgess et al., 2016; Nielsen et al., 2018). To assess the robustness of our community detection results, we have repeated the main analysis after incorporating MGTR into the preprocessing pipeline. To avoid the influence of “artifactually” induced anticorrelations on the community detection, we removed negative values from the correlation matrices prior to the RMT decomposition [note: an example of the non-thresholded MGTR approach can be found in the accompanying Supporting Information (SI) document]. To compare the consistency between the partitions obtained with and without MGTR, we correlated the co-classification matrices obtained from the subject-level clustering of the two methods, and, for interpretability, the percentage of nodes that differ in the subject-derived consensus partitions(i.e., the *Hamming* distance).

In our analysis, we used a multi-modal cortical parcellation (Glasser et al., 2016a) to define the network nodes. To ensure that this did not lead to idiosyncratic results, we have repeated the main analyses with a cortical parcellation by Schaefer et al. (2017) which is based solely on function, and may be more functionally coherent (see SI).

### 2.10. Statistical analysis

Statistical tests for community detection are described above. To compare the similarity measures from the subject-derived vs. group-derived consensus, we used two-tailed permutation hypothesis tests (10,000,000 permutations), and Cohen’s d for the effect size. Fisher’s combined probability test was used to calculate a summary measure of the combined results from the individual permutation tests (at each hierarchical level) bearing on the same hypothesis.

### 2.11. Software and code

The analyses were conducted using MATLAB 2017b (MathWorks Inc., MA, USA). The hierarchical consensus framework was adapted from Jeub et al. (2018) (https://github.com/LJeub/HierarchicalConsensus). We adopted the Louvain implementation from Jeub et al. (2017) (https://github.com/GenLouvain/GenLouvain). The Random Matrix Theory method was adopted from MacMahon and Garlaschelli (2015) (https://mathworks.com/matlabcentral/fileexchange/49011). Miscellaneous network tools were used from the Network Community Toolbox (http://commdetect.weebly.com) and the Brain Connectivity Toolbox (Rubinov and Sporns, 2010) (https://sites.google.com/site/bctnet/Home). The brain plots were vizualized with the Connectome Workbench software (Marcus et al., 2011) (https://github.com/Washington-University/workbench). The code and hierarchical brain maps will be made publicly available once through peer review.

## 3. Results

### Mesoscale organization revealed by subject-derived consensus

We first performed the community detection at the level of the individual subjects’ scans. Using the multi-modal parcellation (MMP) atlas (180 cortical areas per hemisphere, excluding subcortical structures) (Glasser et al., 2016a), regional fMRI time series were used to generate functional networks and corresponding random matrix null models after appropriate preprocessing (see Materials and Methods). These were then used with the Louvain community detection algorithm. To index the full range of spatial resolutions, we applied the algorithm recursively: each daughter community was treated as a new network and the process was repeated until no statistically significant communities were found under the null model. Near-degeneracy of the Louvain algorithm was addressed by considering 100 runs of the algorithm and picking the most stable output before proceeding to the next hierarchical level (see Materials and Methods).

This resulted in a median of 5 hierarchical levels for each subject (range 1-8). The number of communities across all hierarchical levels ranged between 2 and 134. To identify a representative partitioning for the group based on the information from the subject-level partitioning, we adopted a consensus clustering approach (Jeub et al., 2018; Lancichinetti and Fortunato, 2012; Bassett et al., 2013; Sporns and Betzel, 2016). This method consists of summarizing the outputs of the subject clustering results by quantifying the number of times nodes *i* and *j* were assigned to the same partition across subjects and hierarchies and populating a co-classification matrix *C_ij_* with these values. This co-classification matrix was then subjected to a recursive clustering algorithm recently introduced for multiresolution consensus reclustering (Jeub et al., 2018). This resulted in a subject-derived consensus hierarchical tree with 103 levels, ranging from 3 communities at the first hierarchical split to 112 communities at the finest-grained level (Fig. 1a). Thus, the finest branches of the tree contain about 3 nodes (areas).

**Figure 1:**
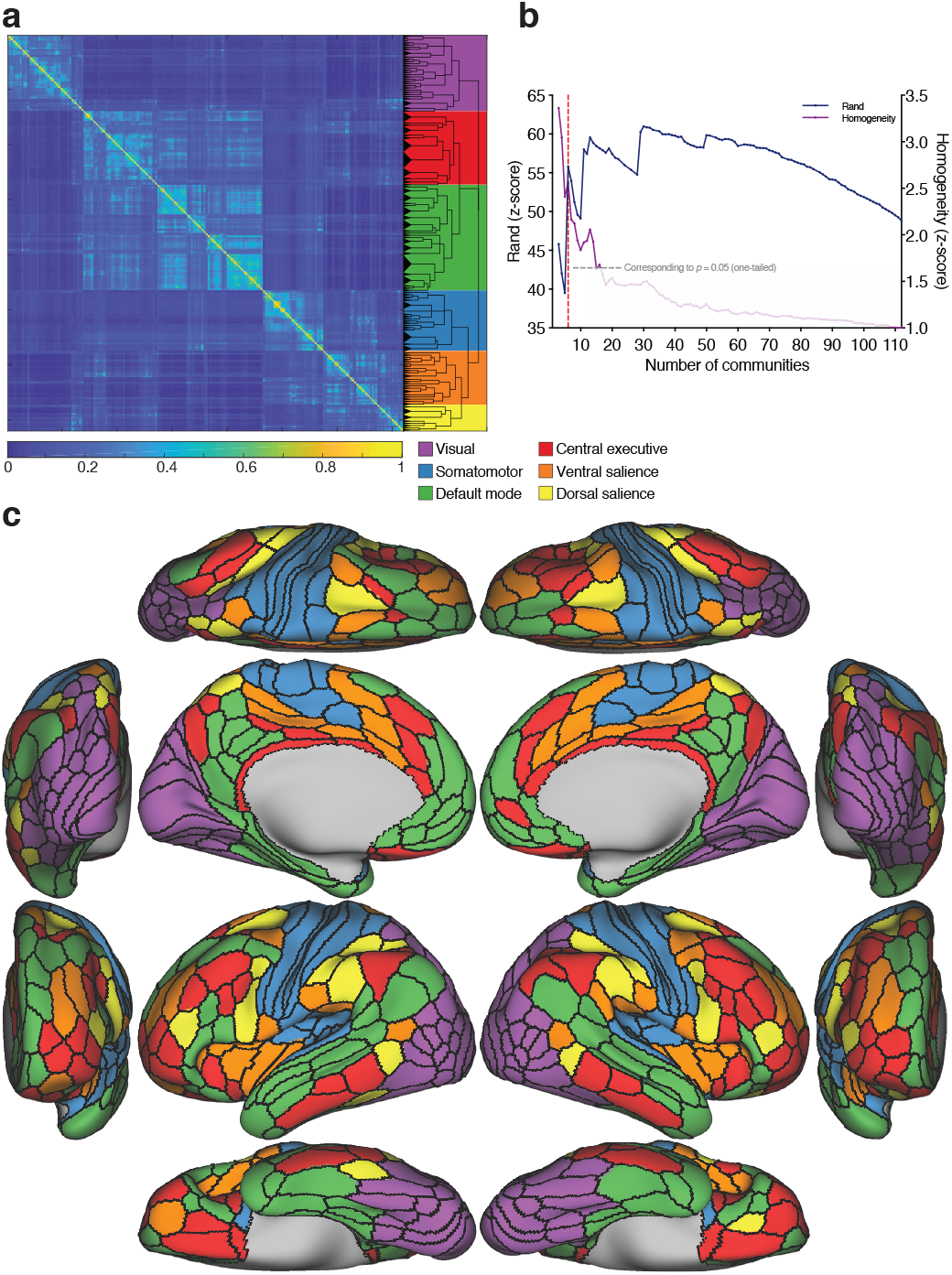
Subject-derived consensus hierarchical partitioning. (a) Co-classification matrix summarizing the results of the subject-level clustering, sorted by community affiliation. The dendrogram represents the hierarchical organization of the nested communities. The length of the arms of the dendrogram are proportional to the average value of the local null model. The background colors represent the candidate division (see below). (b) Similarity plot showing the mean similarity between the partitioning in each hierarchical level in the dendrogram and the clustering at the subject-level quantified by the z-score of the Rand coefficient (blue), and the average z-scored functional homogeneity (purple; values of z > 1.645 represent values that are significantly more homogeneous than the null model at a one-sided *a* < 0.05). The local maximum in similarity corresponds to the partitioning of the cortex into 6 communities (dashed red line). (c) Brain surface plots of the 6 communities corresponding to the local maximum: visual (purple); somatomotor (blue); default mode (green); central executive (red); ventral salience (orange); and dorsal salience (yellow).

By means of the methods that we used, all partition solutions in the hierarchy were statistically significant under the respective null models (see Materials and Methods). To identify partition solutions in the consensus hierarchy that are most expressed at the level of individual subjects, we calculated the average similarity between each consensus partition solution and the subject-level partition solutions (first averaged within each subject across hierarchies, then across subjects). This allowed us to gauge the “representativeness” of the different levels, with those corresponding to local maxima considered to be of particular importance (Fig. 1b).

As an additional independent partition quality metric, we used a measure of functional homogeneity derived from task fMRI. Interestingly, homogeneity peaks appeared to coincide with the similarity peaks, suggesting a convergence between a partition’s “representativeness” and its functional homogeneity across tasks (see SI).

To facilitate the interpretability of the modular organization, here we focus on the first local maximum yielding 6 communities (Fig. 1c). At this level, the organization consisted of an occipital community corresponding to the cortical visual system (*visual*); a community centered around the central fissure and extending to the transverse temporal gyrus, corresponding to the somatosensory, auditory, motor and supplementary cortices (*somatomotor*); a community with anterior and posterior midline components (medial prefrontal and posterior cingulate cortices) as well as a middle temporal component, collectively known as the *default mode*; a community with nodes predominantly in the frontoparietal cortex that is more expressed in the right hemisphere (*central executive*); a cingulo-opercular community spanning the ventral part of the salience system (*ventral salience*); and a more dorsal community spanning dorsal parts of the salience system (*dorsal salience*).

Interestingly, other prominent local maxima appear to have occurred at neurobiologically relevant divisions (Fig. 2). The level yielding 11 communities corresponds to the split of the default mode community into a midline core community and middle temporal lobe community (Fig. 2a). The level resulting in 13 communities represents the split of the auditory community from the somatomotor community (Fig. 2b). The level yielding 19 communities represents the delineation of the language community, more expressed on the left (Fig. 2c). The level yielding 30 communities represents a hemispheric split of the central executive community (Fig. 2d).

**Figure 2:**
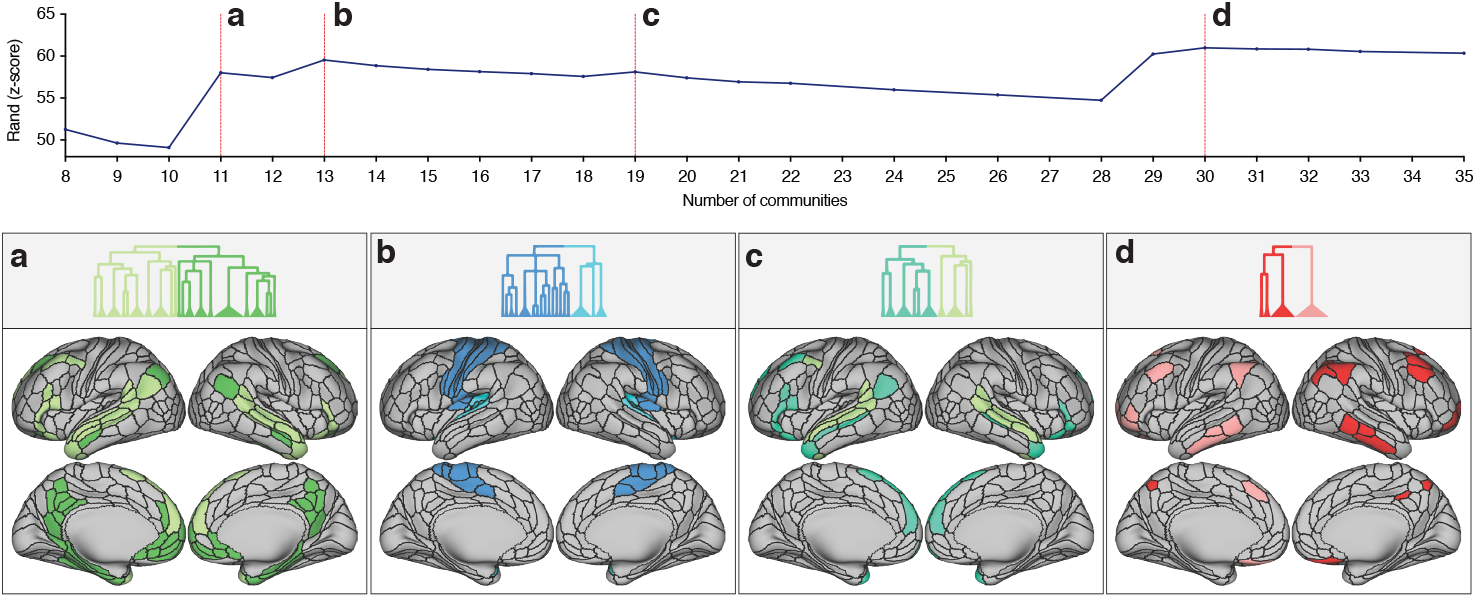
Top: similarity plot showing the mean similarity between the partitioning in each hierarchical level and the clustering at the subject-level quantified by the z-score of the Rand coefficient. The local maxima in similarity are denoted by the dashed red lines. Bottom: emerging communities at local maxima, (a) The level of 11 communities is characterized by splitting of the default mode community into a mainly midline core community (dark green) and mainly middle temporal lobe community (light green), compared to the preceding level. (b) The level of 13 communities is characterized by the splitting of the auditory community (light blue) from the somatomotor community (dark blue). (c) The level of 19 communities is characterized by the emergence of the language community (turquoise) from lateral default mode (light green). (d) The level of 30 communities is characterized by the hemispheric split of the left and right central executive community (red and pink).

### Subject-derived consensus partitions are more accurate representation of individuals ‘ mesoscale features

As a control to the subject-level method, here we perform the community detection on a group-representative network. We concatenated the fMRI regional time series from all participants and generated connectivity and corresponding random matrix null networks. For a meaningful comparison with the subject-level method, we iterated the hierarchical algorithm an equal number of times as the subject-level method (1003 times) and used the multiresolution output from these runs to populate a co-classification matrix. Since the first-level clustering here was performed on a single group-representative network, a slight variability in the outputs of the 1003 runs was deemed necessary to achieve a rich co-classification matrix. For this reason, the procedure to control for the near-degeneracy was applied at the hierarchical consensus clustering only (see Materials and Methods). Each run resulted with a median of 5 hierarchical levels (range 4–5). We then built a co-classification matrix *C_ij_* using the outputs and performed the hierarchical consensus reclustering step. This resulted in a group-derived consensus hierarchy of 101 levels, with a number of communities ranging between 4 and 143 (Fig. 3a). See SI for more details.

**Figure 3:**
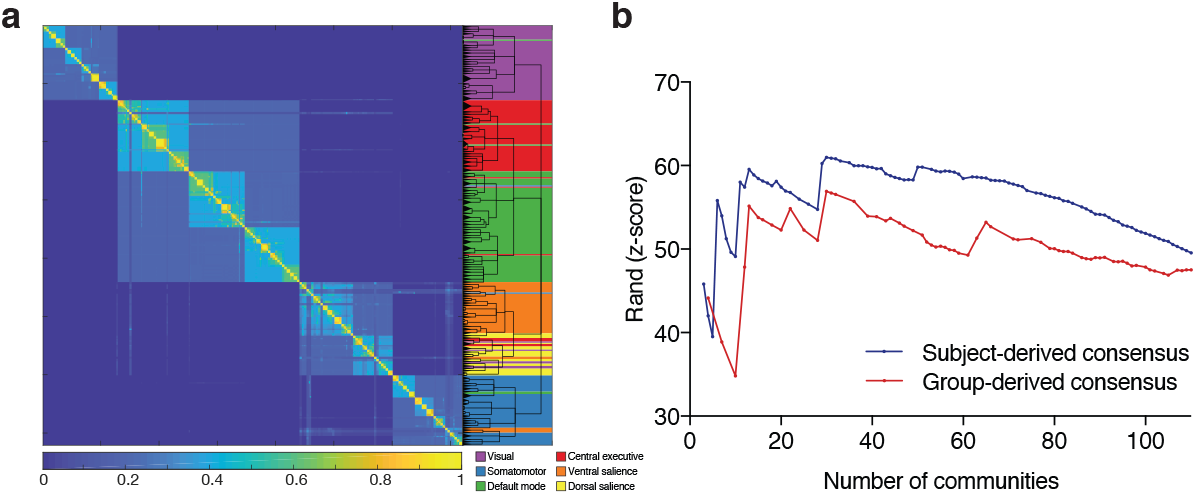
Group-derived consensus hierarchical partitioning. (a) Co-classification matrix summarizing the results of the group-level clustering, sorted by community affiliation. The dendrogram represents the hierarchical organization of the nested communities. The length of the arms of the dendrogram are proportional to the average value of the local null model. The background colors and key are superimposed from the subject-level consensus clustering that yielded 6 modules in Fig. 1a–c. (b) Similarity plot showing the mean similarity between the clustering at the subject-level, and the partitioning in each hierarchical level (number of communities) in the group-derived consensus partitioning (red) and the subject-derived consensus partitioning (blue); similarity is quantified using the *z*-score of the Rand coefficient.

One of the goals of this study was to assess whether applying the clustering algorithms at the subject level followed by meta-reclustering to reach a group consensus (i.e., the method in the previous section) confers a meaningful advantage over applying the clustering algorithm to a group-representative network as is generally done in the literature. For this reason, we calculated the average similarity between the group average-derived consensus partitions (Fig. 3a) and the partitions at the level of the individual subjects. We compared these values with the similarity between the subject-derived consensus partitions and the partitions at the level of the individual subjects (Fig. 3b). Compared to the group-derived consensus partitions, the subject-derived consensus partitions were more similar to the individuals’ partitioning throughout the 65 hierarchical levels that are in common (twotailed permutation tests; *n* = 1003; 10^−7^ < *p* < 2.073 × 10^−4^). The mean effect size was *d* = 0.5381 ± 0.2477 SD (Fisher’s combined probability test, *p* < 10^−16^). One exception is the partitioning that yielded 4 communities, where the group-derived consensus showed higher similarity to individuals’ partitioning two-tailed permutation test; *n* = 1003; *p* = 1.8 × 10^−5^; *d* = 0.1905). We also conducted this analysis with the *normalized mutual information* (Danon et al., 2005) instead of the *z*-score of the Rand coefficient, and obtained similar results (see SI).

### Nodal variability in co-classification consistency partially explained by its topological role

Nodal consistency values ranged between 0.5200 and 0.8817 across hierarchies (mean = 0.7609 ± 0.0582 SD) (Fig. 4a). At the level of 6 communities (Fig. 4b), the multiple regression model that included nodal hubness, nodal signal to noise, and task covariation significantly predicted nodal consistency scores (*R*^2^ = 0.07613, *p* = 1.0337 × 10^−7^, *df* = 356) (Fig. 4b). Nodal hubness was the only term that significantly contributed to the model (standardized estimate = 0.2731, *p* = 1.657 × 10^7^) (Fig. 4c).

**Figure 4:**
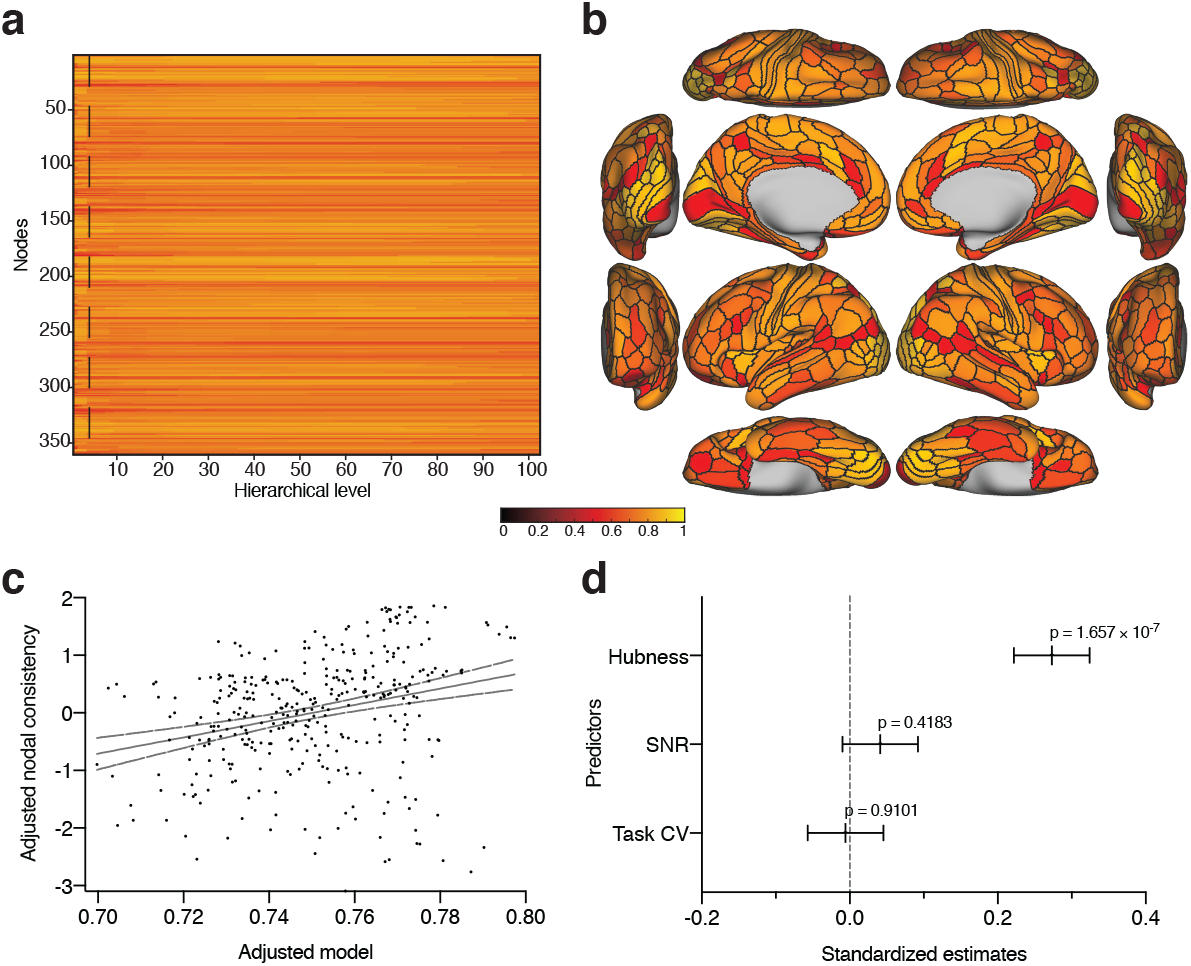
Nodal consistency. (a) Nodal consistency scores across the hierarchies. Dashed line represents the hierarchy that resulted in 6 communities shown in Fig. 1c, and plotted in (b). (c) Multiple regression model that included nodal hubness, nodal SNR, and task coefficient of variation as predictors of nodal consistency. (d) Breakdown of the predictors and their standardized estimates.

### Partitions are robust to prepossessing methods

Subject-derived consensus partitioning after MGTR yielded 115 hierarchical levels, with an organization that is similar to the main analyses (Fig. 5a). The vectorized upper triangles of the co-classification matrix generated from the data with MGTR and without MGTR were strongly correlated (*r* = 0.9538, *p* < 10^−3ω7^, *df* = 64618) (Fig. 5b). Similarly, the partition similarities were high across all hierarchies as evidenced by the z-score of the Rand coefficient (corresponding to *p* < 10^−307^), and the percent agreement ranged between 75.56% and 96.11% (mean = 82.20%, SD = 3.49) for the other hierarchical levels (Fig. 5c).

**Figure 5:**
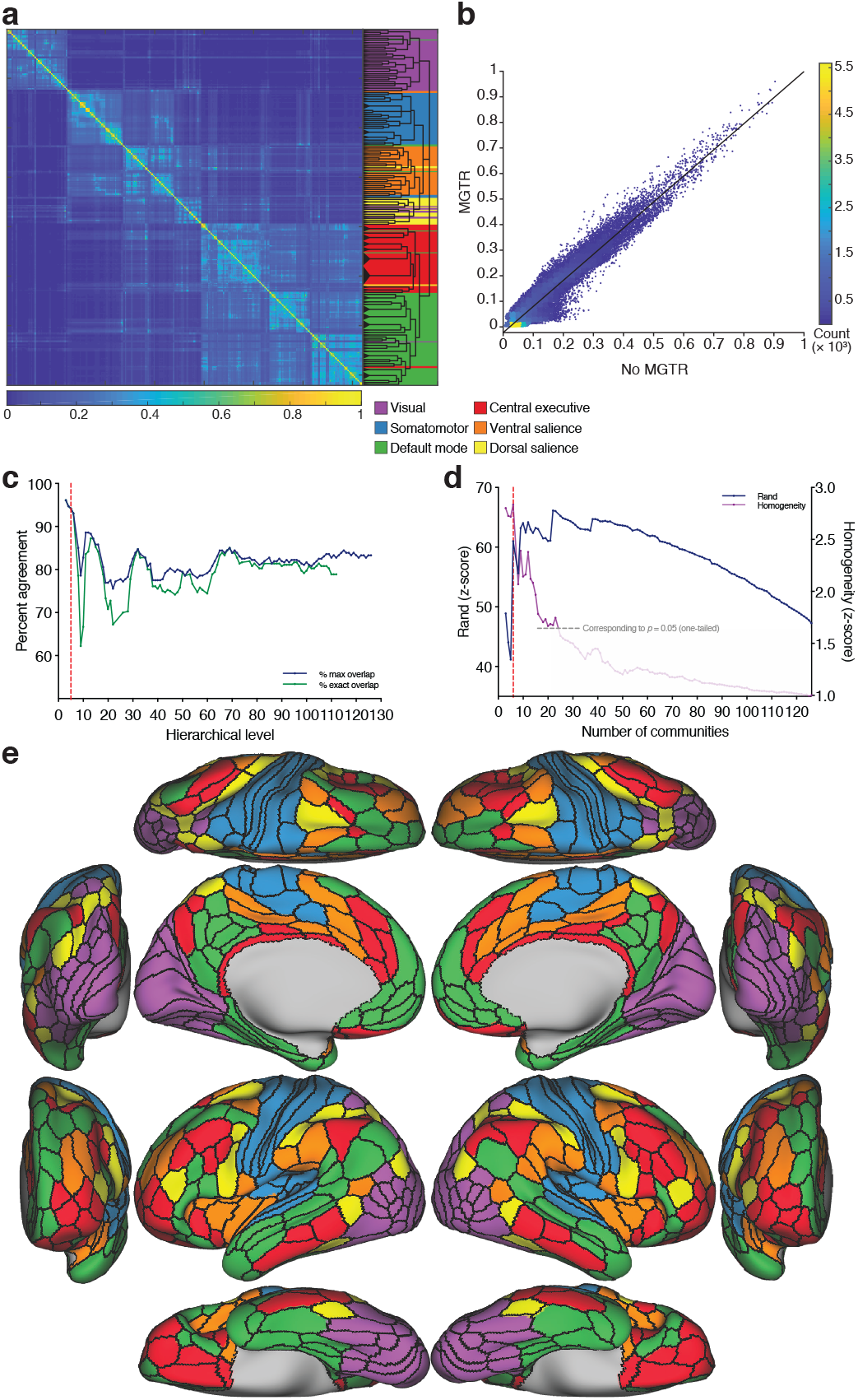
Subject-derived consensus hierarchical partitioning after MGTR. (a) Co-classification matrix and dendrogram. Background colors represent the partitioning from Fig. 1. (b) Scatterplot showing the vectorized entries of the co-classification matrices with and without MGTR. (c) Percent consistency between the subject-derived consensus partitions with and without MGTR. (d) Similarity between the consensus partitioning and the clustering at the subject-level (blue), and the average *z*-scored functional homogeneity (purple; values of z > 1.645 represent values that are significantly more homogeneous than the null model at a one-sided *α* <; 0.05). Similar to Fig. 1, there is local maximum corresponding to the level of 6 communities. (e) Brain surface plots of the 6 communities after MGTR.

As in the main analyses, the similarity of the consensus hierarchies to the subject-level partitioning as well as the homogeneity peaked at the level of 6 communities (Fig. 5d). This partitioning revealed the same subsystems identifid in the main analyses (Fig. 5e), with 93.06% agreement (dashed red line in Fig. 5c).

As an example partition, we also included the partitioning that yielded 22 communities (corresponding to the highest similarity peak) (Fig. 6). At this relatively fine-grained modular organization, there are five visual communities (medial, inferolateral, para, midlateral, superolateral), four somatomotor communities (central, paracentral, inferior, auditory), three ventral salience communities (superior, posterior, insular) two dorsal salience communities (superior, inferior), four central executive communities (right, left, limbic, dorsal anterior cingulate), and four default mode communities (medial, temporal, limbic, language).

**Figure 6:**
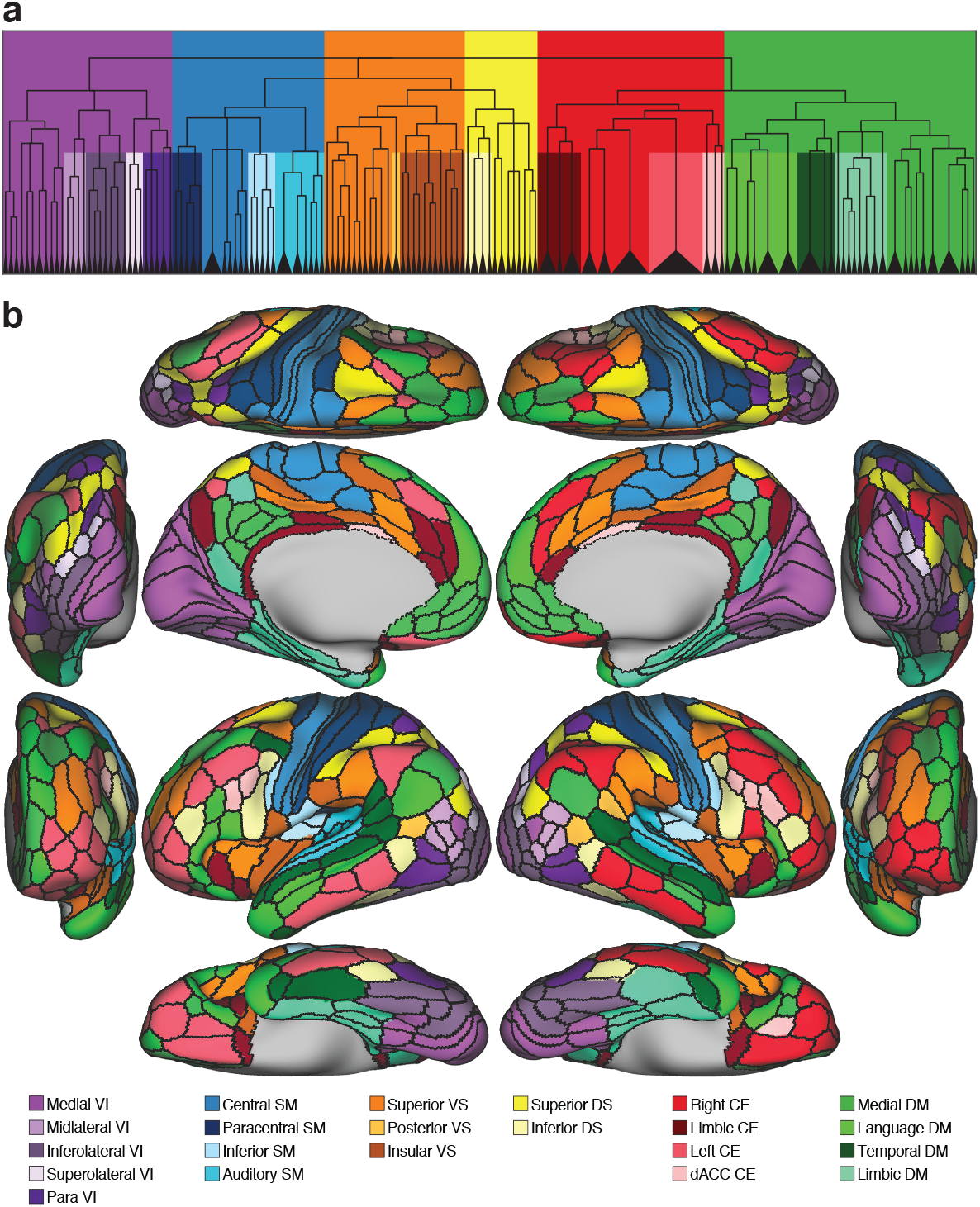
Subject-derived consensus hierarchical partitioning at the level yielding 22 communities. This figure corresponds to the Rand global maximum in Fig. 5d. (a) The dendrogram (from Fig. 5a) and background colors highlighting the fractionation of the 6 main communities into the 22 subcommunities. (b) Brain surface plots of the 22 communities.

## 4. Discussion

Using subject-level clustering of functional networks and a reclustering framework, we mapped the mesoscale architecture of the cortex at multiple scales. Confirming the study hypothesis, the subject-derived consensus framework yielded results that were more similar to the individual subjects compared to the group-derived consensus, despite the fact that both were obtained from the same initial data. While it is inevitable that any type of average will obscure individual features, obtaining a population-level representation that is as faithful as possible to the mesoscale of individuals is preferable. The subject-level consensus enables a representation that better represents a central tendency, resembling individuals to a greater extent. In previous studies, resting-state fMRI acquisitions typically consisted of few minutes (e.g., 5-10 min; TR = 3000 ms), therefore, averaging may have been a good strategy to increase the signal-to-noise ratio. However, in the current study, the acquisitions consisted of 60 min (TR = 720 ms) potentially mitigating this issue.

The implicit assumption that the individual subjects’ clustering results are an appropriate reference is reinforced by a body of literature showing that subject-per-subject extraction of community organization better agrees with the individuals’ behavioral characteristics (Kong et al., 2018). Further, the convergence of the two independent metrics that we used to assess the partition solutions—similarity and task functional homogeneity—gives credence to the notion that the similarity between the consensus partitions and those at the level of the individual is a relevant quality metric (see SI).

Data science approaches are being increasingly implemented to study the relationship between brain connectivity, behavior, and psychopathology. However, a major challenge in this field is the large number of features compared to the number of observations per study. For example, a vertex-wise HCP “dense” connectome includes approximately 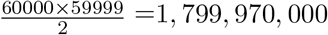 features per subject (unique edges in a symmetrical undirected connectivity network). By using a parcellation atlas (e.g., the neuroanatomically informed multimodal parcellation used in this study) the number of features can be reduced to 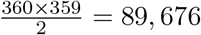. One important aspect of the multiresolution modular atlas that we developed in this study is that it can be used to further reduce the number of features while accounting for the connectome architecture. For example, adopting the partitioning that yielded 22 modules (Fig. 6) would reduce the number of unique features to 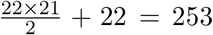, while assessing both inter- and intra-modular connectivity (Akiki et al., 2018).

Several important neuroanatomical observations can be made. The representative partition solution that yielded 6 clusters (level 4; Fig. 1c) corresponds to a well-described set of functional systems (Yeo et al., 2011; Power et al., 2011; Meunier, 2009). The visual system can be seen represented in a large occipital, ranging from early and higher-order visual cortices to the dorsal and ventral streams. Even at the coarsest scale, the visual system comprised a separate community than the somatosensory system. Likely, a parallel of the heavy anatomical differentiation in that area and consistent with electrophysiological studies in humans and primates (Felleman and Essen, 1991). The somatomotor community included both early and higher somatosensory and motor cortices, and the supplementary motor cortex. Relatively early on in the hierarchy, there is a separation of the auditory system (Fig. 2b). The rest of the somatomotor system remains relatively cohesive. The cluster that spans several midline (medial prefrontal cortex and posterior cingulate cortex) and middle temporal structures, is known in the literature as the default mode, and is known to be involved in task-negative processes (Andrews-Hanna et al., 2010; Anticevic et al., 2012). At the level of 6 communities (Fig. 1c), the default mode constitutes a single entity, but subsequently branches into a midline cluster (anterior medial prefrontal and posterior cingulate cortices) and a middle temporal cluster (Fig. 2a). This division is consistent both functionally and anatomically; evidence from task-imaging showing that the midline structures are functionally specialized for self-relevant decisions and the inference of other people’s mental states (i.e., theory of mind), whereas the temporal components are implicated in autobiographical memory (Andrews-Hanna et al., 2010). At the level of 4 communities (Fig. 1a), there is a large “salience” community that is spread over several higher-order associative regions implicated in salience detection and response, which subsequently branches into specialized ventral (also known as the cingulo-opercular system) and dorsal salience systems (Fig. 1c), an organization that has been well described previously (Uddin, 2014). Interestingly, in Fig. 2c, there is branching of a community that is mostly expressed in the left hemisphere and includes Broca’s area and the superior temporal gyrus, and bears resemblance to areas of activation during language processing tasks (Ji et al., 2019; Langs et al., 2015; Glasser et al., 2016a,b). This community, which has traditionally evaded most (Yeo et al., 2011; Power et al., 2011) (but not all (Ji et al., 2019; Langs et al., 2015)) mesoscale investigations, may represent a language system (Ji et al., 2019; Langs et al., 2015). It is notable that this putative language community is substantially more in agreement with the one described in Langs et al. (2015) than Ji et al. (2019), despite the fact that the former used a markedly different analytic approach (voxel-based, clustering in embedding space, etc).

Another interesting observation is the absence of a parallel to the “limbic” network described by Yeo et al. (2011) which was localized to the ventral surface of the brain. A possible explanation is that the older data that was used in their study had marked signal dropout at the base of the brain, where this network was identified. The HCP data used in the current study had significant improvements in terms of a reduction in signal dropout from the ventral brain and other regions due to air sinuses; notably the orbitofrontal, inferior temporal, and lateral mid-temporal cortices (Power et al., 2014b). In our analyses, parcels in this region were assigned to the default mode and the central executive communities. Another possible explanation for this discrepancy could be the varying definition of what a “cluster” means across studies. Here clusters refer to communities, whereas in Yeo et al. (2011), clusters were identified based on connectivity profiles. A recent study by Ji et al. (2019), that used data from the HCP and the MMP cortical parcellation, identified an “oribito-affective” community that corresponds to posterior orbitofrontal parts of the limbic network descibed in Yeo et al. (2011), though the authors note that it had the lowest “confidence score” of community assignment among the identified networks. Much of the nodes in the described orbito-affective community (i.e., posterior orbitofrontal) are included in our central executive community (Fig. 1c).

Disagreements between our community node assignments and those in Ji et al. (2019) could be due to several methodological differences, such as our use of a consensus procedure to avoid a network group average, a multiscale community detection framework, and a null model based on RMT for community detection that is more compatible with functional networks (compared to the more prevalent permutation null models). Because functional networks are estimated using correlation measures, they are subject to indirect effects that manifest themselves as artifactual connections. These indirect effects are due to second-, third- and higher-order interactions that are not present in the real (i.e., *direct*) network [for example, if nodes 1 and 2 and nodes 1 and 3 are strongly correlated, a correlation will also be observed between node 2 and 3 even where none exists (or, if a true relationship exists, here it would be overestimated)] (Zalesky et al., 2012; Barzel and Barabási, 2013). This often leads to an inaccurate estimation of the network modular structure, if not accounted for by the null model (Bazzi et al., 2016; MacMahon and Garlaschelli, 2015). The RMT-based community detection method that we employed mitigates these effects. In a recent study, a permutation null model failed to recover the ground truth community from neurophysiological recordings in the suprachiasmatic nucleus in mice, while an RMT-based null model was successful (Almog et al., 2017). Another potential contributor to the discrepancy is the choice of the spatial scale of interest: while the current study characterized a wide range of spatial scales, and opted for a data-driven method to focus on those that are most expressed at the level of individuals in addition to functional homogeneity, Ji et al. (2019) described the organization at the scale yielding 12 clusters, based on a priori “hard criteria” (e.g., a requirement for a separate auditory community), in addition to stability metrics. It is therefore conceivable that exploring different spatial resolutions in their data could show more consistent parallels.

Calculating the nodal consistency revealed that certain nodes were less consistent than others in terms of community assignments (Fig. 4). To ensure that this was not caused by signal dropout in these particular regions, we calculated the SNR from the time course. This analysis did not reveal an association between the nodal SNR and consistency scores. Another potential reason that we wanted to rule out was that low consistency nodes are those that are highly task-specific. To quantify that, we used the group-wise task fMRI activation map contrasts and calculated a measure of how much variability each node is exhibiting in terms of activation across different tasks. That also did not turn out to contribute to the variance of the nodal consistency scores. The other potential explanation was that these nodes play a specific—hub-like—topological role, integrating information from different modules, and because their connectivity profiles is not primarily limited to a single module, they are less consistently classified. While there are numerous metrics that index hubness (Rubinov and Sporns, 2010), we opted for the simplest definition—nodal strength. This hubness metric was found to partially account for the variability in consistency (contributing approximately 7.6% of the variance). Although it only explained a small part of the variance, it remains entirely possible that the nodal strength is insufficient to capture hubness. In fact, indexing hubness in correlation networks is a known challenge and traditional metrics do not rigorously identify important hubs (Power et al., 2013). In fact, one of the alternate definitions proposed in Power et al. (2013), which is based on finding nodes in which multiple modules are represented, is conceptually similar to our consistency metric.

Processing the data with and without MGTR resulted in broadly similar partitions (Fig. 5). We were able to say that the subject-derived consensus partitions resemble individual-level partitions more than the group-derived consensus because both were derived from the same data. However, the same framework could not be used to compare consensus partitions derived with and without MGTR as the comparator is not the same. Despite the fact that MGTR removes some relevant physiological neural data present in the global signal and may alter the properties of the networks especially the negative edges [which we had to remove (see SI)], it is very effective at removing motion-related artifact (Power et al., 2017a,b). Evidence that supports “sacrificing” some neural data in favor of a less contaminated signal includes a recent observation that MGTR strengthens the associations between connectivity and behavioral measures (Li et al., 2019). Therefore, we release all atlases without taking a stance regarding which method is superior.

Since the MMP atlas that we adopted is limited to the cerebral cortex, our analysis did not comprise the subcortex or cerebellum. However, we recognize the importance of whole-brain the mapping, and to inform the literature, have repeated the main analysis after adding subcortical (Fan et al., 2016) and cerebellar nodes (Diedrichsen et al., 2009) from other atlases (see SI). Some studies have assigned community affiliations for the subcortex or cerebellum on a post hoc basis after identifying the cortical clusters (Yeo et al., 2011; Buckner et al., 2011; Choi et al., 2012; Ji et al., 2019). To avoid “imposing” the identified cortical modules on the other structures, we have included all nodes at the same time in the clustering algorithm. The cortical systems remained largely consistent, and the sub-cortical and cerebellar nodes were predominantly assigned to the default mode, central executive, and a salience modules, with only a few nodes assigned to the visual and somatomotor modules (see SI). The results did not fall in line with the classical neuroanatomical literature (Kandel et al., 2012) (e.g., striatal structures being part of a somatomotor modules or thalamic nuclei wither their respective cortical systems). This could be due to several reasons. While cortical communities are largely assortative (Betzel et al., 2018), subcortical and cerebellar communities may require a broader definition of communities that include non-assortative types (e.g., core-periphery and disassortative communitites) (Betzel et al., 2018). Therefore, future whole-brain endeavors should test methods that allow for this greater diversity (e.g., stochastic block modeling (Betzel et al., 2018), overlapping communities (de Reus et al., 2014), and approaches that allow for a varying spatial configuration of functional brain regions (Bijsterbosch et al., 2018), and temporal dynamics (Bassett et al., 2013)).

While we used a multimodal gradient-based parcellation because it is believed to be the most neuroanatomically accurate to date (Glasser et al., 2016a), other atlases based on unimodal analyses of functional connectivity may be more functionally-coherent (Schaefer et al., 2017; Gordon et al., 2016; Craddock et al., 2011). We repeated the main analysis using the Schaefer et al. (2017) atlas and recovered a broadly similar set of modules (see SI). However, a quantitative analysis (Alexander-Bloch et al., 2018) was not within the scope of this article, but should be pursued in the future. In this study, we constrained the analysis to the cerebral cortex, though we recognize the importance of extending the mapping of these communities. to the subcortex and cerebellum (Fan et al., 2016; Diedrichsen et al., 2009). The multimodal brain atlas that we adopted does not include any subcortical or cerebellar nodes. Some groups have added community affiliations for the subcortex or cerebellum on a post hoc basis (Buckner et al., 2011; Choi et al., 2012; Ji et al., 2019). The challenge in our case is to keep the definition of a community consistent. This will likely require the treatment of the nodes of the whole brain at the same time. The decision to define nodes as brain parcels instead of a voxel-wise analysis was motivated by factors that include using neurobiologically-meaningful building blocks, mitigating MRI-related limitations, and computational intractability (Eickhoff et al., 2018). However, voxel-wise investigations (e.g., Braga and Buckner (2017); Gordon et al. (2017b)) are equally important in brain mapping endeavors as they naturally provide a finer level of granularity. Finally, other mesoscale organizations should also be explored, such methods include stochastic block modeling (Betzel et al., 2018), overlapping communities (de Reus et al., 2014), and approaches that allow for varying spatial configuration of functional brain regions (Bijsterbosch et al., 2018).

## Disclosure

C.G.A. has served as a consultant or on advisory boards for Genentech, Janssen, Lundbeck, and FSV7, serves as editor for the journal *Chronic Stress* published by SAGE Publications, Inc, and filed a patent for using mTOR inhibitors to augment the effects of antidepressants (filed on Aug 20, 2018).

## Acknowledgements

Data were provided by the Human Connectome Project, WU-Minn Consortium (Principal Investigators: David Van Essen and Kamil Ugurbil; 1 U54MH091657) funded by the 16 NIH Institutes and Centers that support the NIH Blueprint for Neuroscience Research; and by the McDonnell Center for Systems Neuroscience at Washington University. Funding support was provided by the U.S. Department of Veterans Affairs National Center for PTSD and NIH (MH-101498). We would like to thank Samaneh Nemati for data curation. We acknowledge the computational resources provided by Yale University and the National Center for PTSD.

