## Supporting Information for "Determining the Hierarchical Architecture of the Human Brain Using Subject-Level Clustering of Functional Networks"

---

---

### 1. A note on the $z$-score of the Rand coefficient

Comparing the similarity between partition solutions with a markedly different number of communities poses a significant challenge in clustering analyses. It is expected that as the consensus-derived partitions become more fine-grained (a large number of small communities), the calculated similarity with the subject-level partitions will be higher, simply as a function of the number and size of modules (i.e., even if there is no true change in similarity). Calculating the $z$-scores of the Rand coefficient accounts for this bias, and the resulting score can be interpreted as the similarity between two partitions, beyond what can be explained by the number and size of partitions. Another advantage of using the standardized variant of the Rand index is that while there are a number of indices that can be used to measure partition similarity (e.g., normalized mutual information, Jaccard, Minkowski, Adjusted Rand), it has been shown that their $z$-scores coincide (Traud et al., 2011; Fortunato and Hric, 2016).

However, to ensure the robustness of the conclusion that the subject-derived consensus is more similar to the individual compared to the group-derived consensus, we repeated the analysis with the normalized mutual information—another commonly used measure of clustering similarity (Danon et al., 2005). The analysis with the normalized mutual information yielded the same conclusion as with the $z$-score of the Rand coefficient that we describe in the main text: overall, compared to the group-derived consensus partitions, the subject-derived consensus partitions had a higher mean similarity to the individuals' partitioning (Fig. S1). The mean effect size across all 65 levels that are in common was $d = 0.1514 \pm 0.06807$ SD (Fisher's combined probability test, $p < 10^{-16}$). Individually, of the

*Corresponding author

65 levels, 56 were statistically significant (two-tailed permutation tests; $n = 1003$; $10^{-7} < p < 0.05$). All but one level had a higher mean similarity for the subject-derived consensus. The one exception is the partitioning that yields 4 communities, where the group-derived consensus had a non-statistically significant higher similarity (two-tailed permutation test; $n = 1003$; $p = 0.3062$; $d = 0.0461$).

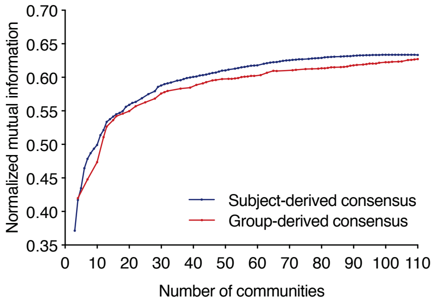


Figure S1: Similarity plot showing the mean normalized mutual information (similarity) between the clustering at the subject-level, and the partitioning in each hierarchical level (sorted by number of communities) in the group-derived consensus partitioning (red) and the subject-derived consensus partitioning (blue).

### 2. A note on the relationship between functional homogeneity and subject similarity

The primary data-driven metric that we used to select which partition solutions in the hierarchical tree to focus on—quantified by the Rand coefficient—was based on the premise that partitions that are highly expressed at the level of individual subjects were particularly salient. As an additional metric that is independent, we also calculated a measure of *functional homogeneity* derived from task fMRI contrasts (see Materials and Methods in the main text).

Qualitatively, it was noteworthy that the extrema of the Rand and of the homogeneity metric were often co-occurring (i.e., Fig. 1b and Fig. 5b in the main text, and Fig. S4). To quantify this agreement, we started by selecting partitions solutions that were homogeneous above chance (one-sided $\alpha < 0.05$, which translates into $z$-score $> 1.645$). Next, to focus on the peaks, we removed the linear trend from the Rand and the homogeneity vectors using the `detrend.m` function in MATLAB (Fig. S2b). We found a positive correlation between the detrended homogeneity and similarity scores (Fig. S2c; Combined data points: $r = 0.6022$, $p = 2.4507 \times 10^{-5}$, $n = 42$; Subject-derived: $r = 0.8381$, $p = 3.4676 \times 10^{-4}$, $n = 13$; Group-derived: $r = 0.8653$, $p = 0.0012$, $n = 10$; Subject-derived with MGTR: $r = 0.3134$, $p = 0.1913$, $n = 19$). The convergence of the two independent metrics was reassuring, and supports that they may be surrogates for partition quality.

However, a comparison of the methods (e.g., subject-derived vs. subject-derived with MGTR vs. group-derived) based on homogeneity was not conducted because the task fMRI activation maps that we used in our calculations were from a group average, and so statistical testing could not be adequately performed. However, even if the analysis was repeated on all participants to enable statistical testing, the difference in $z$-scores that we have observed with the task fMRI average map were small (Fig. S2a), and even if there is a "statistically significant" difference in homogeneity between the groups, the magnitude of the effect would not be meaningful on a practical level.

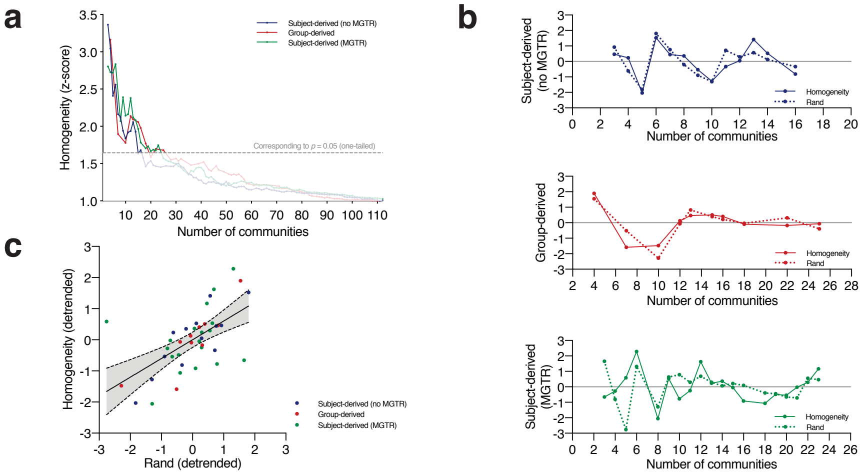


Figure S2: Subject-derived consensus hierarchical partitioning after MGTR. (a) Co-classification matrix and dendrogram summarizing the results after MGTR and thresholding. Background colors represent the partitioning from Fig. 1. (b) Scatterplot showing the vectorized entries of the co-classification matrices. (c) Percent and Rand consistency between the MGTR partitioning and the original partitioning.

### 3. A note on the regression of the mean grey-matter "global" time course regression

Global signal regression (e.g., MGTR) has a long contentious history but it has been shown to be one of the most effective preprocessing steps to reduce the effects of motion (Power et al., 2017, 2014), letting true patterns in the data emerge (Li et al., 2019; Nielsen et al., 2018). Negative edges in particular are known to be controversial (Murphy et al., 2009; Fox et al., 2009; Power et al., 2014). While in the original analysis without MGTR we included the full range of correlations (positive and negative), we had to exclude negative values after MGTR, as not doing so resulted in aberrant results. For example, no separation between default mode and visual communities—two very distinct and well-defined communities (Vincent et al., 2007; Yeo et al., 2011), despite fine-graned fractionation of other communities (Fig. S3).

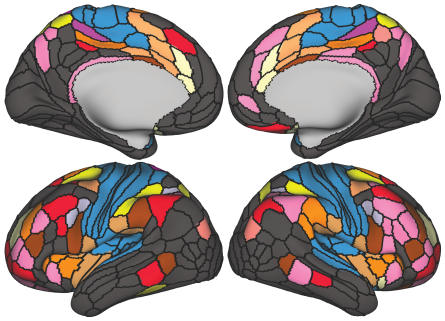

Figure S3: Example partition with MGTR without excluding negative edges. Not excluding negative edges after MGTR leads to aberrant results. Notice how the large community in dark grey did not split into distinct default mode and the visual communities, despite fine-grained splitting of other communities (other colors).

### 4. Results from the group-derived consensus

The communities look broadly similar to the subject-derived consensus, with certain exceptions. One difference is that there is no similarity peak before the level of 13 communities (Fig. S4a). In fact, there is absence of a 6-community solution (the closest being a 7-community solution with the additional module consisting of a subcommunity in the somatomotor community).

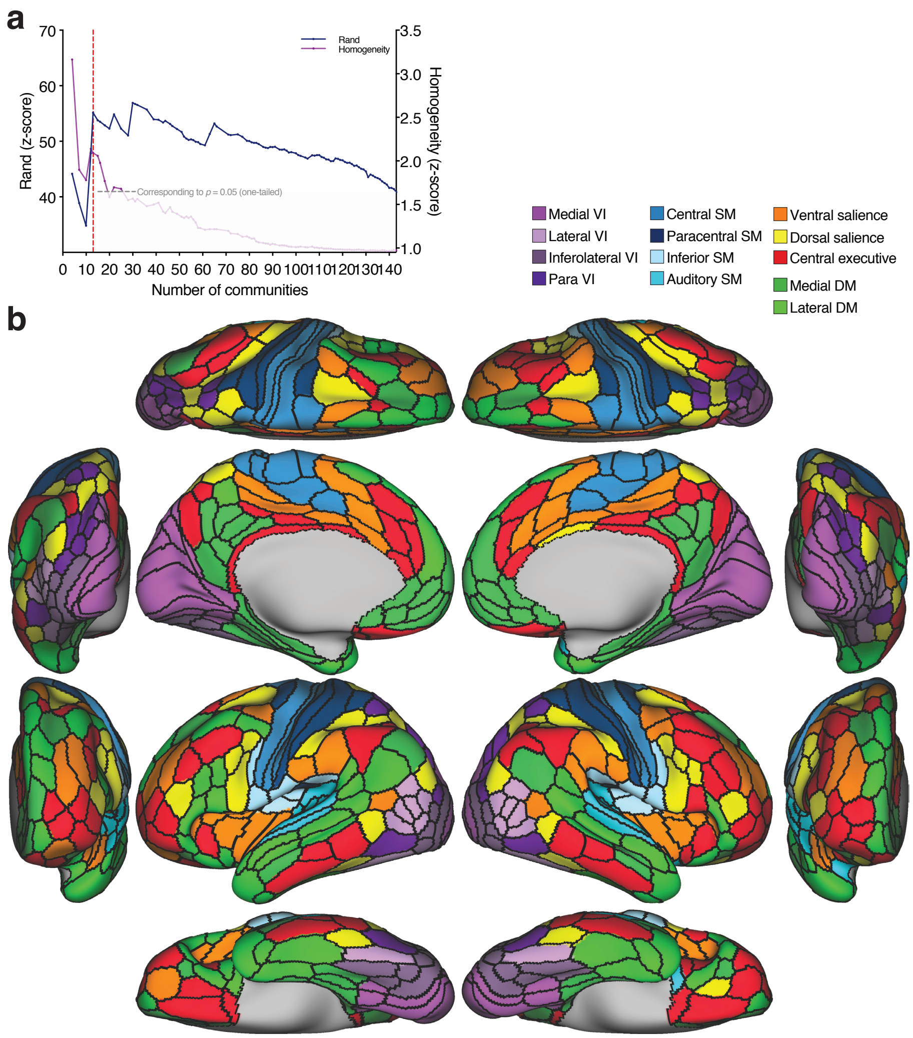


Figure S4: Group-derived consensus hierarchical partitioning. (a) Similarity between the consensus partitioning and the clustering at the subject-level (blue), and the average $z$-scored functional homogeneity (purple; values of $z > 1.645$ represent values that are significantly more homogeneous than the null model at a one-sided $\alpha < 0.05$). There is local maximum corresponding to the level of 13 communities (dashed red line). (b) Brain surface plots of the 13 communities.

### 5. Parcellation with an alternate functional atlas of the cerebral cortex

For the main analysis we used the multi-modal parcellation (MMP) because it is thought to be the most neuroanatomically informed to date (Glasser et al., 2016). However, since the parcels were not solely based on function, it is possible that the nodes may be less "functionally coherent" compared to a parcellation that is solely based on function (Eickhoff et al., 2018; Schaefer et al., 2017). To ensure that the results of the 6-community partition in the main text were not an idiosyncratic result of the parcellation atlas, we repeated the analysis using a recently developed functional parcellation of the cortex (Schaefer et al., 2017). Under this scheme, the cortex is divided into 400 regions of interest.

Using the same methodology, we obtained a subject-derived consensus (Fig. S5a). Similar to the results of the MMP parcellation reported in the main text, the $z$-score of the Rand coefficient showed a local maximum the level of 6 communities (Fig. S5b). At this level, the two parcellation atlases appear show a similar mesoscale organization (Fig. S5c). Certain dissimilarities are also notable. For example, certain regions with the primary and early visual cortex were not assigned to the visual community, possibly suggesting an integrative function (Tong, 2003). In contrast, in the MMP atlas, this region was parcellated differently (as it incorporated anatomical information as well), making the average time course from that parcel more coupled to neighboring visual areas. Although there are methods to quantify the agreement between different atlases (Alexander-Bloch et al., 2018), such a comparison is not within the scope of this current article, but should be pursued in the future.

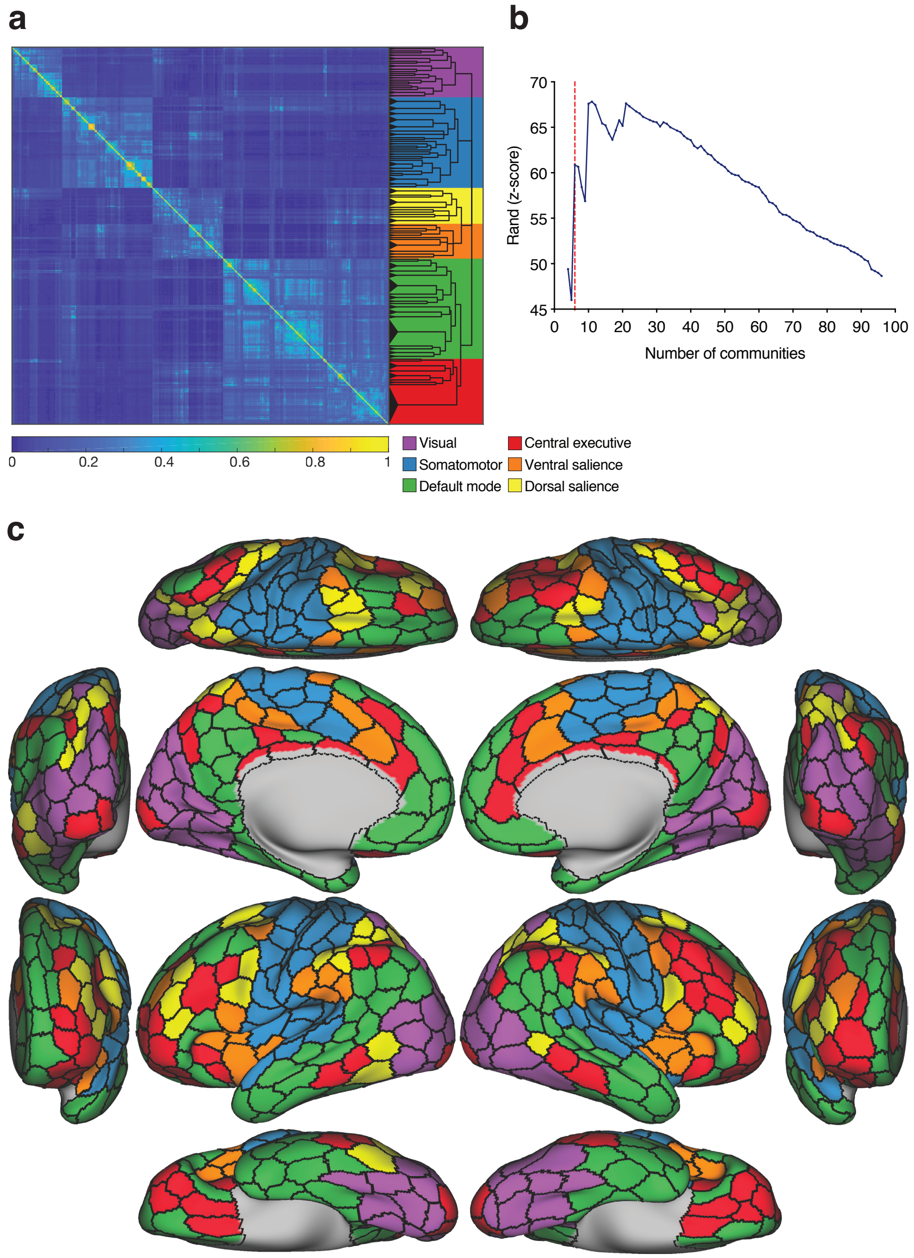


Figure S5: Subject-derived consensus hierarchical partitioning with the Schaefer parcellation atlas. (a) Co-classification matrix summarizing the results of the subject-level clustering, sorted by community affiliation. The dendrogram represents the hierarchical organization of the nested communities. The length of the arms of the dendrogram are proportional to the average value of the local null model. The background colors represent the candidate division (see below). (b) Similarity plot showing the mean similarity between the partitioning in each hierarchical level in the dendrogram and the clustering at the subject-level quantified by the $z$-score of the Rand coefficient. The local maximum in similarity corresponds to the partitioning of the cortex into 6 communities (dashed red line). (c) Brain surface plots of the 6 communities corresponding to the local maximum.
